# JNK signalling regulates self-renewal of proliferative urine-derived renal progenitor cells via inhibition of ferroptosis

**DOI:** 10.1101/2022.08.24.505101

**Authors:** Lisa Nguyen, Michelle Westerhoff, Leonie Thewes, Wasco Wruck, Andreas S. Reichert, Carsten Berndt, James Adjaye

## Abstract

With a global increase in chronic kidney disease patients, alternatives to dialysis and organ transplantation are needed. Stem cell-based therapies could be one possibility to treat chronic kidney disease. Here, we used multipotent urine-derived renal progenitor cells (UdRPCs) to study nephrogenesis. UdRPCs treated with the JNK inhibitor-AEG3482, displayed decreased proliferation and downregulated transcription of cell cycle-associated genes as well as the kidney progenitor markers -SIX2, CITED1, and SALL1. In addition, levels of activated SMAD2/3, which is associated with the maintenance of self-renewal in UdRPCs, were decreased. JNK inhibition resulted in less efficient oxidative phosphorylation and more lipid peroxidation via ferroptosis-an iron-dependent non-apoptotic cell death pathway linked to various forms of kidney disease. Our study reveals the importance of JNK signalling in maintaining self-renewal as well as protection against ferroptosis in SIX2-positive UdRPCs. We propose that UdRPCs can be used for modelling ferroptosis-induced kidney diseases.

## Introduction

Due to the increasing numbers of patient mortality due to kidney-associated diseases, new medical options besides the conventional dialysis and organ transplantation are needed. However, shortage of donor organs and immune compatibility restrict these therapeutic approaches. A future therapy option may be cell replacement therapies with pluripotent stem cells (PSCs)-derived cellular products; however, these have drawbacks that include ethical concerns and tumorigenicity. Counter measurements to the use of PSCs are the cost-effective and non-invasive urine cells. Cell types isolated from urine include urine stem cells, podocytes, and proximal tubule epithelial cells (Oliveira Arcolino et al., 2015). Highly interesting for research and medical purposes is the subpopulation of multipotent urine stem cells (USCs) which display typical MSC characteristics (Pavathuparambil Abdul Manaph et al., 2018; Rahman et al., 2020; Sato et al., 2019; Zhang et al., 2014). Besides expressing kidney-specific markers such as Podocin and Synaptopodin (Pavathuparambil Abdul Manaph et al., 2018; Sato et al., 2019), USCs express the renal progenitor marker sine oculis homeobox homolog 2 (SIX2). Based on SIX2 expression we conveniently coined the term urine-derived renal progenitor cells (UdRPCs) (Rahman et al., 2020). Presumably, UdRPCs originate from the upper urinary tract of the kidney, which was indicated by the presence of Y-Chromosome in these cells, isolated from the urine of a female patient who received a kidney transplant from a male donor (Bharadwaj et al., 2013).

The proliferative potential of UdRPCs and their ability to differentiate into various kidney cell types, such as podocytes (Erichsen et al., 2022) and tubular cells (unpublished) make them a promising tool for kidney regeneration experiments. Results from our earlier study, suggested the maintenance of self-renewal of UdRPCs via FGF and TGFβ signalling with modulations on JNK signalling (Rahman et al., 2020). JNK is part of the mitogen-activated protein kinase (MAPK) family and each pathway is a sequential activation of MAPKKKs (Javelaud and Mauviel, 2005). This pathway is involved in various processes such as proliferation, survival as well as stress and inflammation-induced apoptosis and is also linked to acute and chronic kidney diseases (Awazu, 2017; Engel et al., 1999; Smith et al., 2021; Tournier et al., 2000). In nephron progenitor cells JNK signalling plays a major role in maintaining the progenitor fate.

In the last decades, the cellular process of ferroptosis was discovered to play a role in the emergence of kidney diseases (Tang and Xiao, 2020). Ferroptosis was first described by Dixon *et al*. in 2012 as an iron-dependent form of non-apoptotic cell death. It is characterized by accumulation of iron that leads in combination with peroxides to lipid peroxidation (Hu et al., 2019; Li et al., 2020). Glutathione peroxidase 4 (GPX4) catalyzes the glutathione (GSH)-dependent reduction of H2O2 and hydroperoxides of polyunsaturated fatty acids to its corresponding alcohols, thus restricting the cellular levels of peroxides and repairing oxidized lipids (Galluzzi et al., 2018; Ingold et al., 2018). A depletion of glutathione levels and a consequent decrease in GPX4 activity leads to the accumulation of lipid peroxidation and subsequently to membrane destruction (Galluzzi et al., 2018; Jiang et al., 2021). Other circumstances as activated mitochondrial voltage-dependent anion channels and mitogen-activated protein kinases, stress on endoplasmic reticuli, and inhibited cystine/glutamate antiporter may also induce ferroptosis (Jiang et al., 2021; Li et al., 2020). Ferroptosis-induced morphology changes include blistered cell membrane, lack of chromatin condensation, less and more fragmented mitochondria with denser membranes and reduction or disappearance of cristae (Galluzzi et al., 2018; Li et al., 2020). Thereby, ferroptosis is involved in various diseases of the brain, heart, liver, and kidney (Conrad et al., 2021; Galluzzi et al., 2018). Studies on mice demonstrated a lack of fully functional GPX4, a key suppressor of ferroptosis, resulted in acute kidney failure (Friedmann Angeli et al., 2014; Tonnus et al., 2021). Besides acute kidney failure, ferroptosis-induced diseases in the kidney include acute kidney injury, I/R injury, Clear Cell Renal Cell Carcinoma and Adrenocortical Carcinomas (Galluzzi et al., 2018).

For this study, we investigated the effects of JNK signalling on three different UdRPC cultures by applying the inhibitory compound AEG3482 for 72 h. We were able to connect JNK signalling with the maintenance of proliferation and self-renewal in UdRPCs. Moreover, JNK inhibition induced ferroptosis and reduced cellular oxidative phosphorylation via disruption of the mitochondrial membrane potential. We propose that UdRPCs treated with a JNK inhibitor can be used for modelling ferroptosis-induced kidney diseases and promotion of JNK signalling in UdRPCs may represent a novel strategy for kidney regeneration experiments.

## Results

### High concentrations of JNK inhibitor induce cell death

First, optimal concentrations of the JNK inhibitor AEG3482 were tested on UM51 cells. Tested concentrations were 10 µM, 50 µM and 100 µM. The treatment was conducted for 2 to 5 days with daily medium change. The UM51 cells tolerated a concentration of 10 µM without any morphological changes and were cultivated for up to 5 days before stopping the treatment due to high cell confluency (Figure S1A). A concentration of 50 µM led to a morphology change from the typical rice grain-like to a more flattened appearance with partial detachment and indistinctive cell membrane (Figure S1A). The highest dose of 100 µM caused complete cell detachment and eventually cell death after 48 h (Figure S1A). UF21 cells treated with 50 µM and 100 µM JNK inhibitor could be kept for 72 h before the cells died (Figure S1B). As the concentrations of 50 µM and 100 µM were highly stressful to the cells, therefore, a concentration of 10 µM AEG3482 was used for all following experiments.

### JNK inhibition leads to decreased proliferation and loss of progenitor state

Cell morphology changes in UM51, UM27 and UF21 cells after JNK inhibition with AEG3482 were observed after 72 h Figure 1A). In all three cell cultures, JNK inhibition resulted in lower confluency compared to their respective controls. The proliferation capacity of the UdRPCs with and without JNK inhibition at 72 h was analysed by immunofluorescence-based staining for KI67 (Figure 1B). Reduced numbers of KI67^+^-proliferative cells were observed under JNK inhibition, which was confirmed by a proliferation assay displaying the ratio of Ki67^+^ cells against the total cell number (Figure 1*B-C*). An unpaired t-test analysis confirmed a highly significant downregulation of KI67^+^ UM51 cells (p < 0.001) and significant downregulation in UM27 cells (p < 0.05), while there was no significant difference between control and JNK inhibition in UF21 cells (Figure 1*C*). mRNA expression of progenitor- and cell cycle-related genes was analysed via semi-quantitative real-time PCR. The progenitor markers SIX2, SALL1 and VCAM1 as well as KI67 were downregulated in JNK-inhibited UdRPCs (Figure 1D). A heatmap depicted the expression of *CCND2, SMAD4, CDC14B, HDAC1, CCNH, WEE1, TFDP2* and *RBX1* in the controls, but down-regulated in JNK-inhibited UM51 cells in the time points 24 h, 72 h and 120 h Figure 1E).

**Figure 1.**
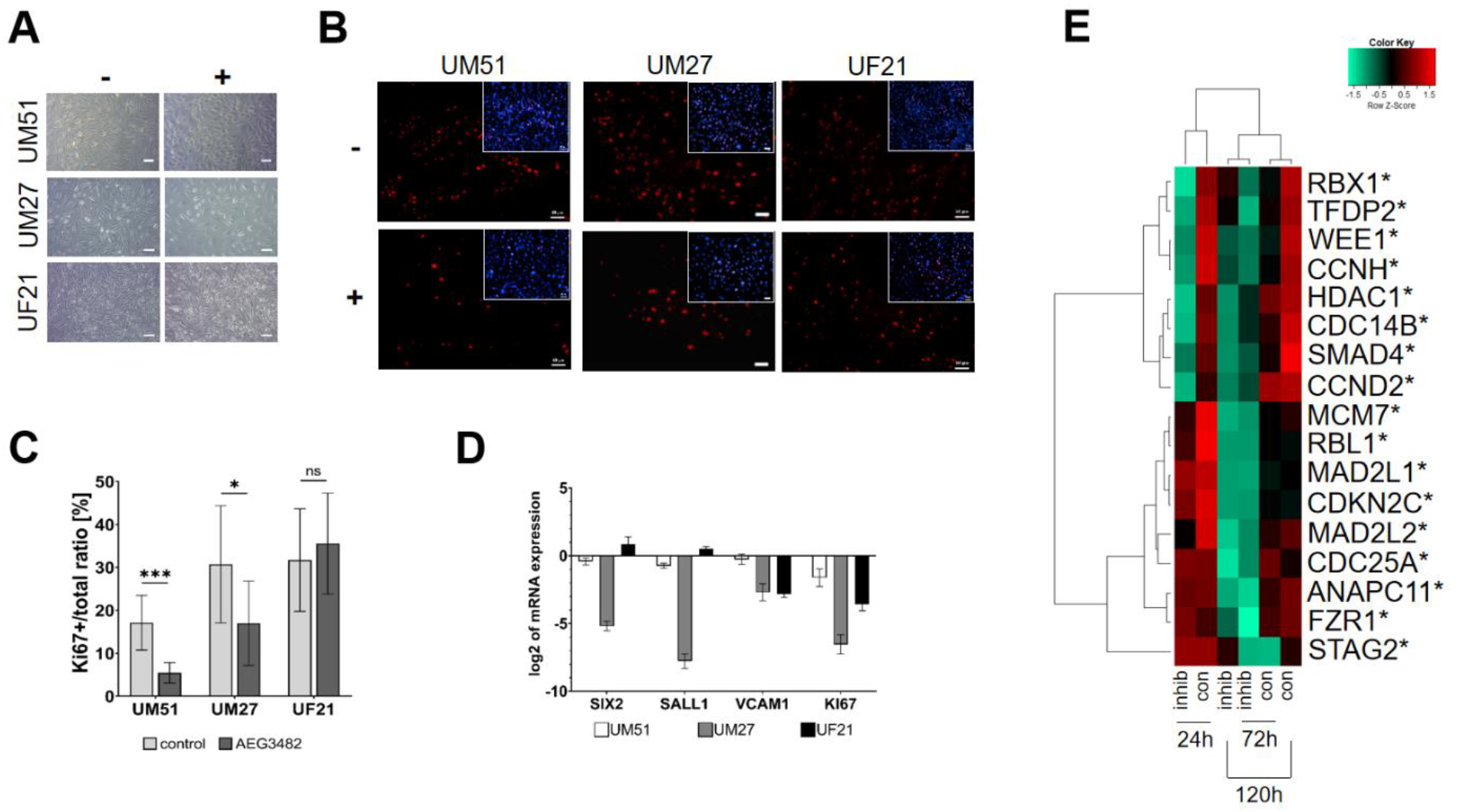
Inhibition of JNK reduces the proliferation of UdRPCs. A) Morphology of the three UdRPCs UM51, UM27 and UF21 with or without AEG3482 treatment after 72 h. Scale bars represent 100 µm. B) Ki67 expression in UdRPCs treated with or without AEG3482 for 72 h. Scale bars represent 50 µm. C) Ki67 proliferation assay for JNK-inhibited UdRPCs (n=10; * ρ-value < 0.05, ** ρ-value < 0.01, *** ρ-value < 0.001). D) mRNA expression of nephron progenitor marker SIX2, SALL1, VCAM1 and Ki67. Mean values were normalized to the housekeeping gene RPL37A. Error bars indicate SEM. E) Gene expression of cell cycle-related genes for the time points 24 h, 72 h and 120 h depicted in a Pearson’s heatmap.

### Cell cycle-related processes are regulated by JNK inhibition

Microarray-based gene expression profiles of UM51 cells treated with JNK inhibitor for adjusted time span and their specific controls were analysed via transcriptome analysis. A comparison of gene expression between control and JNK inhibition were displayed in Venn diagrams for the time points 24 h, 72 h and 120 h (det p >0.05). Green colour represents the control, while red denotes JNK inhibition (Figure 2A-C). Complete gene lists for all time points 24 h, 72 h and 120 h are listed in Table S2-4. Kyoto Encyclopedia of Genes and Genomes (KEGG) pathway analysis was performed using exclusive gene-sets pertinent to control and AEG3482 treatment. Significantly downregulated in all three time points (24 h, 72 h, 120 h) was the KEGG pathway cell cycle (ratio >1.5) (Table S5). We were able to pinpoint 6 genes common in all three time points, which we found to be regulated by JNK *(BUB1, CCNA2, CCNB2, CCND2, MCM7, PLK1)* (Table S5).

**Figure 2.**
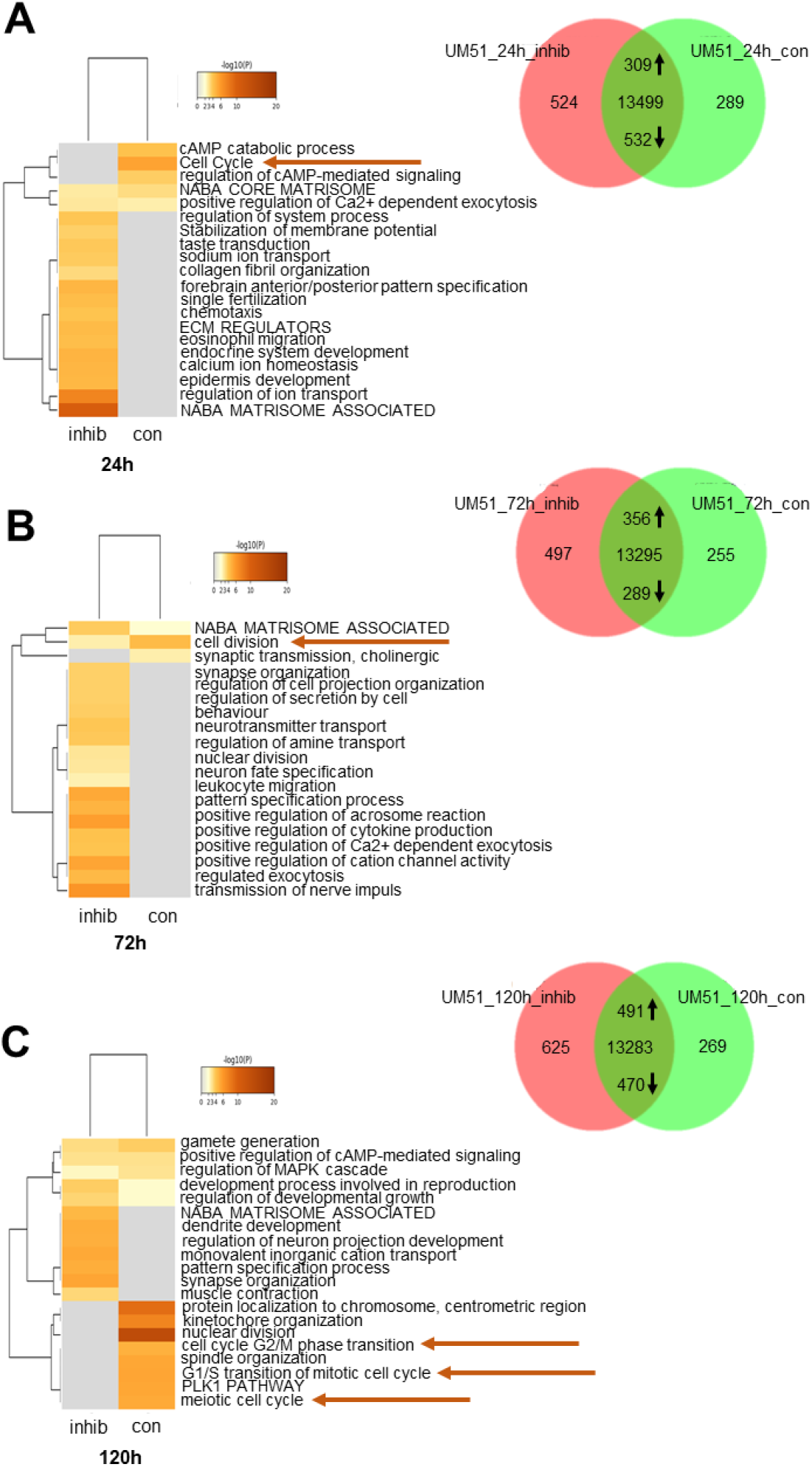
JNK signaling is associated with cell cycle processes in UdRPCs. A) Representative enrichment clusters for control and JNK-inhibition after 24 h depicted in a heatmap and cell cycle-related processes marked with an arrow. Venn diagram of control and JNK-inhibited UM51 cells for the time point 24 h. B) Representative enrichment clusters for control and JNK-inhibition after 72 h depicted in a heatmap and cell cycle-related processes marked with an arrow. Venn diagram of control and JNK-inhibited UM51 cells for the time point 72 h. C) Representative enrichment clusters for control and JNK-inhibition after 120 h depicted in a heatmap and cell cycle-related processes marked with an arrow. Venn diagram of control and JNK-inhibited UM51 cells for the time point 120 h.

Additionally, Metascape-generated enrichment analyses were processed based on the exclusive gene sets in JNK inhibition and control of all three time points (0.3 kappa score). Representative terms of enrichment clusters related to cell cycle and cell division (Table S6) with the highest P-values in the UM51 data sets of each time point were represented by heatmaps (Figure 2A-C). Notably, the GO BP terms related to cell cycle lacked enrichment in the JNK inhibitions of 24 h, 72 h and 120 h compared to high enrichment in the specific controls (marked with an arrow in (Figure 2A-C).

### The downstream target phospho-cJUN is downregulated by JNK inhibition

Next, expression of JNK and downstream target cJUN and its phosphorylated form before and after JNK inhibition was examined (Figure 3A-B). Successful inhibition of JNK phosphorylation by JNK inhibitor AEG3482 was confirmed in all three cell cultures (Figure 3A). Interestingly, the level of p-JNK in treated UF21 cells was less reduced than in the other two cell lines (Figure 3A). The protein level of non-phosphorylated t-JNK was not affected by JNK inhibition (Figure 3A). JNK inhibition resulted in a reduced phosphorylation of downstream target cJUN in UM51 and UM27, but not in UF21 cells (Figure 3B). Immunofluorescence-based analysis revealed a lack of differential expression of cJUN and phosphorylated cJUN in untreated and treated UdRPCs (Figure 3C).

**Figure 3.**
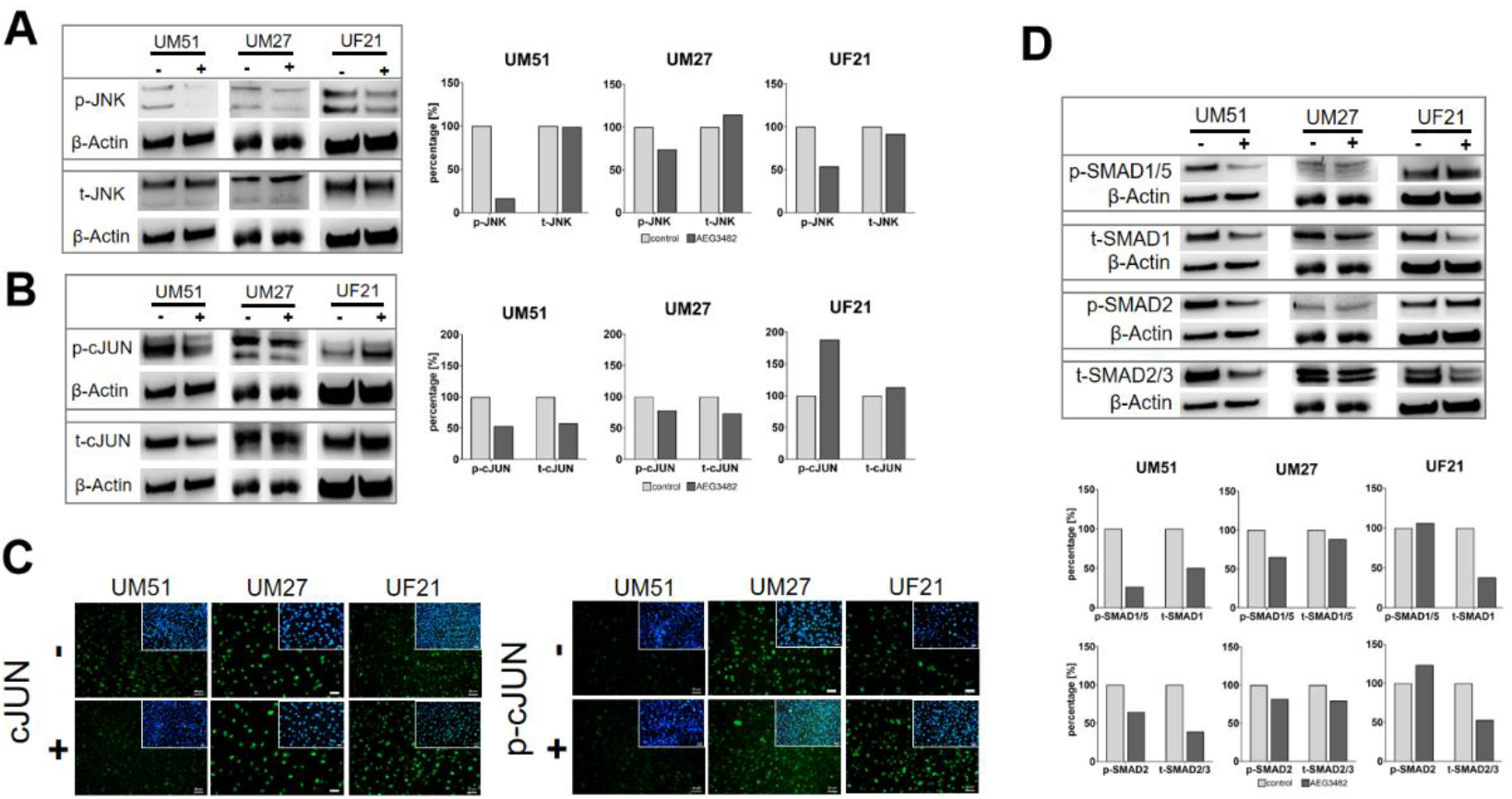
Inhibition of JNK signaling affects the downstream target cJUN and SMAD proteins. A) Protein expression of p- and t-JNK in UdRPCs with or without JNK inhibition. B) Protein expression of p- and t-cJUN in UdRPCs with or without JNK inihibition. C) Immunofluorescence stainings of cJUN and p-cJUN in UdRPCs with or without JNK inhibition. Scale bars represent 50 µm. D) Protein expression of p-/t-SMAD1/5 and p-/t-SMAD2/3 in UdRPCs with or without JNK inhibition.

As our previous study demonstrated the relevance of TGFβ-signalling in the maintenance of self-renewal, protein expression of SMAD2/3 and SMAD1/5 under influence of JNK inhibition was studied. Interestingly, the level of phosphorylated SMAD1/5 and SMAD2 decreased after JNK inhibition in UM51 and UM27 but not in UF21 cells (Figure 3D). Protein levels of total-SMAD2/3 and total SMAD1 were reduced in all three cell cultures (Figure 3D).

### UdRPCs are more susceptible to ferroptosis upon JNK inhibition

KEGG pathway analysis revealed significant upregulation of numerous KEGG pathways including ferroptosis and glutathione metabolism in JNK-inhibited UM51 cells at all time points (ratio <0.67) (Table S5B). Following the results of the KEGG analysis, we decided to carry out an in-depth analysis of the ferroptosis pathway. Since lipid peroxidation is a major hallmark of ferroptotic cell death, the accumulation of lipid peroxides was measured in JNK-inhibited UdRPCs (Figure 4A). The cells exhibited a significant increase in lipid peroxidation, indicating increased sensitivity to ferroptosis. Treatment with the ferroptosis inhibitor Liproxstatin-1 protected the cells from lipid peroxidation (Figure S1C). Additionally, we could also observe a significant increase in lipid peroxidation when FGF signalling was inhibited (Figure S1D). JNK inhibition led to increased expression of gene sets involved in iron and glutathione metabolism (Figure 4B-C). Interestingly, protein expression of the key mediator of the removal of lipid peroxides, GPX4, did not show any difference between control and JNK-inhibition (Figure 4D). Analysis of the mRNA expression of genes related to the glutathione and iron metabolism, confirmed the findings of the heatmap analyses, as the genes *GLC* and *GLCM* and *HMOX1, SLC11A2* and *TFR1* were upregulated upon treatment (Figure 4E-F).

**Figure 4.**
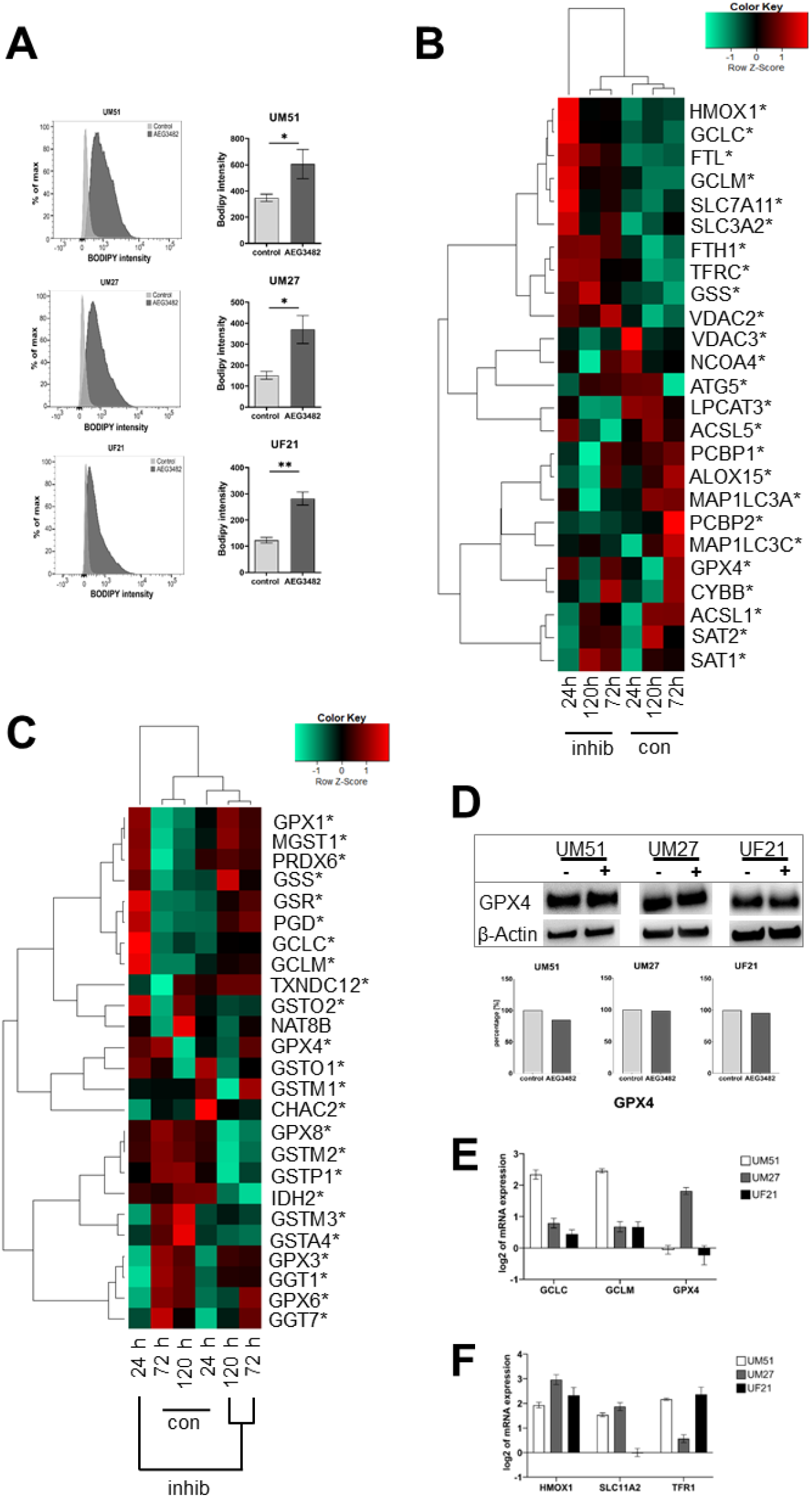
Inhibition of JNK signalling increases lipid peroxidation. A) Representative histograms of measured fluorescence intensities after BODIPY staining and the respective bar plots of mean fluorescence intensity of control or JNK-inhibited UdRPCs (n=5; * ρ-value < 0.05, ** ρ-value < 0.01, *** ρ-value < 0.001). Error bars indicate SEM. B) Gene expression of iron metabolism-related genes in UM51 cells for the time points 24 h, 72 h and 120 h depicted in a Pearson’s heatmap. C) Pearson’s heatmap depicting gene expression of glutathione metabolism-related genes in UM51 cells for the time points 24 h, 72 h and 120 h. D) Protein expression of GPX4 in UdRPCs with or without JNK inhibition. E) mRNA expression of glutathione metabolism-related markers GCLC, GCLM, GPX4. Error bars indicate SEM. F) mRNA expression of iron metabolism-related markers HMOX1, SLC11A2 and TFR1. Mean values were normalized to the housekeeping gene RPL37A. Error bars indicate SEM.

### JNK inhibition partially disrupts the mitochondrial membrane potential and reduces respiration

Oxidative Phosphorylation in JNK-inhibited and control UdRPCs was evaluated by measuring the oxygen consumption rate (OCR) via Seahorse Agilent Mito Stress test. The graphs in Figure 5A show basal respiration, the maximal respiratory capacity and the spare respiratory capacity in UdRPCs treated with the JNK inhibitor. The data obtained demonstrate a significant reduction in basal respiration, maximal respiration and spare respiratory capacity in UM51, UM27 and UF21 cells (Figure 5A).

**Figure 5.**
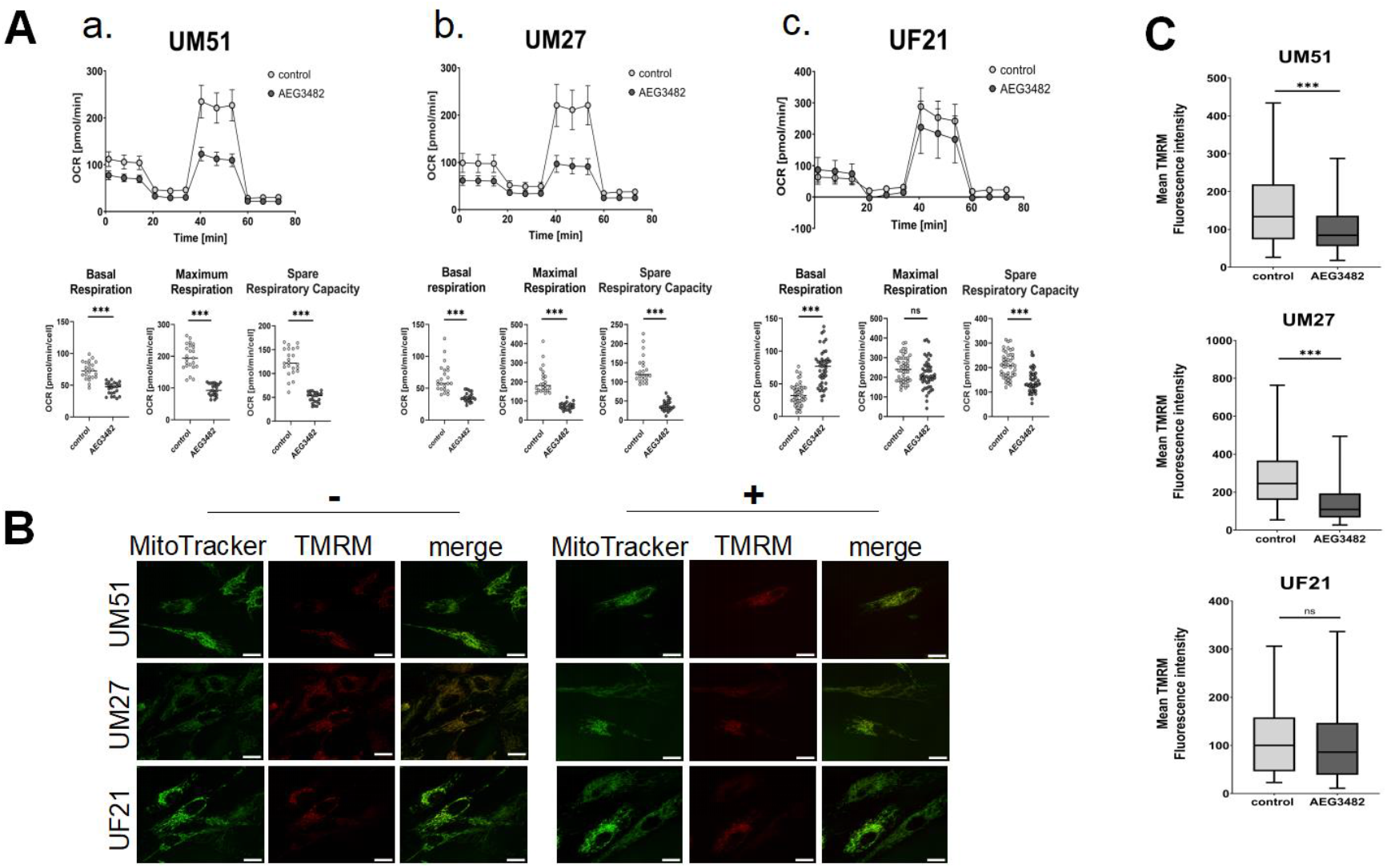
JNK signaling regulates mitochondrial respiration via membrane potential. A) Measurement of OCR in real time in UdRPCs. Basal respiration, maximal respiration and spare respiratory capacity are depicted (* ρ-value < 0.05, ** ρ-value < 0.01, *** ρ-value < 0.001). Error bars indicate SD. **a**. OCR in UM51 (n=1; N=23). **b**. OCR in UM27 (n=1; N=23). **c**. OCR in UF21 (n= 1; N= 46). B) Fluorescence images of MitoTracker Green and TMRM stainings in UdRPCs UM51, UM27 and UF21 with and without JNK inhibition. Scale bars depict 10 µm. C) Measurement of mean TMRM fluorescence signal intensity in UdRPCs * ρ-value < 0.05, ** ρ-value < 0.01, *** ρ-value < 0.001). Outliers were removed. Error bars indicate SEM. **a**. Mean TMRM signal in UM51 is depicted (n=2; ctrl. N= 1; + JNK inhibitor N= 2). **b**. Mean TMRM signal in UM27 is depicted (n=2; ctrl. N= 1; + JNK inhibitor N= 2). Error bars indicate SEM. **c**. Mean TMRM signal in UF21 is depicted (n=2; ctrl. N= 1; + JNK inhibitor N= 2).

Reduction of oxidative phosphorylation and SRC are indicators of defective mitochondrial function. Therefore, we investigated the influence of JNK inhibition on mitochondrial membrane potential in UdRPCs via TMRM staining (Figure 5B). We observed a significant reduction of mitochondrial membrane potential in UM51 and UM27 compared to the respective controls (Figure 5C). UF21 cells showed no change in mitochondrial membrane potential upon JNK inhibition (Figure 5C).

## Discussion

### JNK inhibition decreases the proliferation and leads to loss of progenitor character

In this study, JNK signalling was inhibited using the small molecule inhibitor AEG3482 on three UdRPC cultures, UM51, UM27 and UF21. We discovered the importance of JNK for the proliferative potential of the self-renewing urine-derived renal progenitor pool.

Different concentrations of the inhibitor were tested beforehand, and the final concentration of 10 µM AEG3482 was used further. Inhibition of the active JNK enzyme was demonstrated by Western Blot analysis. The inhibitor did not completely block the phosphorylation of JNK, which would explain why the active form of the direct downstream target cJUN was still observed (Javelaud and Mauviel, 2005). Interestingly, in UF21 cells, an inhibitor concentration of 10 µM resulted in higher p-cJUN protein levels of treated than in non-treated cells. Besides the observation that UF21 cells could tolerate higher concentrations of the JNK inhibitor AEG3482 for longer periods, adaptation or insensitivity to the 10 µM concentration of the inhibitor might be an explanation for higher p-cJUN levels.

Even though the cell morphology of the three UdRPC cultures did not change upon JNK inhibition, we observed lower cell density. We therefore assumed a reduced proliferation rate in the AEG3482-treated cells. Comparable results were found in a study on the role of JNK signalling in proliferative nephron progenitors of a mouse model (Blank et al., 2009). While it was demonstrated that BMP7 signalling enhanced the proliferation of murine SIX2^+^ nephron progenitor cells, JNK inhibition disrupted this effect (Blank et al., 2009). It has been shown that BMP7-induced JNK signalling increases the proliferative capacity of SIX2^+^ nephron progenitor cells in the mouse model. As UdRPCs share similar characteristics with nephron progenitor cells, a similar outcome regarding the proliferation is very likely. To test the effect of JNK inhibition on the proliferative capacity of UdRPCs, we conducted a follow-up proliferation assay using KI67 as marker. We could detect statistically significant reduction of proliferation in UdRPCs, UM51 and UM27, treated with JNK inhibitor after 72 h. The proliferation rate of JNK-inhibited UF21 cells was not significantly reduced, which might be caused by higher inhibitor tolerance as exhibited before. Moreover, gene expression of KI67 and gene ontologies related to cell cycle were downregulated in JNK-inhibited UdRPCs. In our study, genes of KEGG annotated cell cycle-related GO term (*BUB1, CCNA2, CCNB2, CCND2, MCM7, PLK1)* were downregulated by JNK inhibition. The identified common genes are mainly involved in the cell cycle phase transitions, indicating interruption of the cell cycle and subsequent reduction of proliferation. Additionally, a heatmap analysis revealed absence of cell cycle-related genes (*CCND2, SMAD4, CDC14B, HDAC1, CCNH, WEE1, TFDP2, RBX1*) upon JNK inhibition and expression in all controls as well as sustained expression in JNK-inhibited UM51 after 24 h. We observed a downregulation of the progenitor marker *SIX2, SALL1* and *VCAM1* (*CD106*) after AEG3482 treatment. Besides the effect of JNK on the proliferative capacity of UdRPCs, reduced expression of the genes *SIX2, SALL1* and *CD106* indicate a loss of progenitor status. Muthukrishnan *et al*. confirmed the Tak1-Jnk-Jun pathway maintained the numbers murine nephron progenitors by keeping the cells in a proliferative state (Muthukrishnan et al., 2015). Therefore, our results signify that JNK signalling is involved in cell cycle progression, proliferation and maintenance of the progenitor state of UdPRCs.

*In vivo*, NPC self-renewal is sustained by the interaction of growth factors, signalling pathways and metabolic pathways within the nephron progenitor niche and changes can induce differentiation (Liu et al., 2017; Oxburgh and Rosen, 2017). In our previous work, we have observed that FGF-induced TGFβ/BMP signalling determines the cell fate of urine-derived renal progenitor cells, since we have demonstrated downregulation of p-SMAD2/3 and upregulation of p-SMAD1/5/8 in differentiated UdRPCs upon activation of WNT signalling (Rahman et al., 2020). We hypothesized that active SMAD2 is necessary to maintain the progenitor state of UdRPCs. In contrast to our previous findings, we observed downregulation of both, p-SMAD2 and p-SMAD1/5, upon JNK inhibition. This indicates an interconnection between JNK and SMAD signalling, which is unexpected as SMAD signalling is commonly believed to be independent from JNK signalling (Zhang, 2009). Based on our findings from previous works (Rahman et al., 2020), the decreased expression of p-SMAD2/3 implies the loss of nephron progenitor state due to JNK inhibition. Other signalling pathways such as the BMP7-induced and JNK-independent Smad1/5/8 pathway were also observed to be contributing to the maintenance of the renal progenitor pool in the murine model (Tomita et al., 2013). Tomita *et al*. (2013) were able to demonstrate that an inhibition of BMP7-Smad signalling leads to the differentiation of nephron progenitor cells and thus they proposed an important role of BMP7-Smad signalling for the maintenance of the renal progenitor cells and the determination of final nephron numbers (Tomita et al., 2013). Similar to the mentioned observations in the mouse model, phosphorylated SMAD1/5/8 was decreased in JNK-inhibited UdRPCs UM51 and UM27, indicating the loss of the progenitor state.

### Active JNK signalling protects urine-derived renal progenitor cells against ferroptosis

In this study, we have shown that the inhibition of JNK induces ferroptosis. Ferroptosis is characterized by the iron-dependent formation of lipid peroxides leading to cell death. The FACS-based measurement of lipid peroxides confirmed that JNK inhibition significantly increased sensitivity to ferroptosis. Moreover, addition of a ferroptosis inhibitor (Liproxstatin-1) protected the cells from accumulating lipid peroxides. FGF signaling regulates the maintenance of self-renewal in NPCs and UdRPCs (Barak et al., 2012; Rahman et al., 2020). Inhibition of FGF signaling via FGFR inhibitor SU-5402 increased the level of lipid peroxides significantly. A recent publication described the role of FGF21 in the suppression of iron overload-induced ferroptosis in liver (Wu et al., 2021). Similarly, FGF signaling may have protective properties against ferroptosis in UdRPCs, however this warrants further investigation beyond the scope of the current study.

KEGG pathway analysis revealed a significant upregulation of genes associated with the GO term ferroptosis and glutathione metabolism upon JNK inhibition. In our heatmap analyses, we observed clustered expression of ferroptosis-related genes in the JNK-inhibited samples, which are mainly involved in iron and glutathione metabolism. In line with these results, we demonstrated upregulated mRNA expression of glutathione metabolism-related genes *GCLC, GCLM*, and *SLC7A11* (*cystine-glutamate antiporter Xc-*) as well as iron metabolism-related genes *HMOX1, SLC11A2* and *TFR1*. Increased expression of regulators of the iron metabolism involved in the iron (Fe^2+^) release from cellular storage and influx of iron can be one of the inducers of ferroptosis (Chen et al., 2020). An accumulation of free iron (Fe^2+^) catalyzes the Fenton reaction, which generates hydroxyl radicals, thus leading to peroxidation of polyunsaturated fatty acids (Latunde-Dada, 2017). Heme oxygenase 1 (HO-1, encoded by *HMOX-1*) could contribute to increased iron levels by liberation of iron during heme degradation (Kwon et al., 2015). In contrast, other publications describe that HO-1 suppresses ferroptosis. Adedoyin *et al*. demonstrated increased expression of HO-1 in renal proximal tubular cells resulted in alleviation of ferroptosis (Adedoyin et al., 2018). Moreover, HO-1 is a marker for active NRF2, a transcription factor activating the expression of several genes encoding anti-ferroptotic proteins, including GPX4 (Nishizawa et al., 2022).

Interestingly, we could not observe a change in the protein level of GPX4 post JNK inhibition. However, upregulated transcript levels of genes encoding proteins important for GSH biosynthesis indicate that the cell is limited in GSH, the essential cofactor for GPX4 activity. High levels of GSH and increased amounts of GPX4 are usually negative regulators of ferroptosis (Sharma et al., 2021), while the opposite induces this form of cell death (Berndt and Lillig, 2017). Ferroptosis-was discovered to be one of the causes in the induction of kidney diseases such as acute kidney injury, I/R injury, Clear Cell Renal Cell Carcinoma and Adrenocortical Carcinomas (Galluzzi et al., 2018). Therefore, we assume that UdRPCs treated with the JNK inhibitor AEG3482 may represent an easily available model for studying ferroptosis-induced kidney diseases in the near future.

### JNK signaling is involved in the metabolic activity associated with the maintenance of self-renewal in UdRPCs

The role of mitochondria in ferroptosis is not understood so far (Jiang et al., 2021). Mitochondria could promote ferroptosis via formation of peroxides by altered electron transfer during oxidative phosphorylation but could also be a target of ferroptosis by inducing damage to mitochondrial membranes leading also to dysfunctional energy metabolism. Since JNK inhibition is accompanied by ferroptosis, we decided to study the respiratory function of mitochondria in JNK-inhibited UdRPCs. Therefore, Mito Stress test was performed to determine the influence of JNK inhibition on mitochondrial respiration in UdRPCs. We observed a significant reduction of mitochondrial respiration in UdRPCs upon JNK inhibition. In particular, measurement of the spare respiratory capacity (SRC) is an indicator of cellular health, since it represents the cell’s ability to react to increased energy demand or stress. Therefore, significant reduction of the SRC in all three UdRPCs cultures demonstrates a loss of metabolic capacity of mitochondria upon inhibition of JNK signaling.

Oxidative phosphorylation is crucially dependent on the membrane potential generated by the electron transport chain (ETC) in the inner mitochondrial membrane (Perry et al., 2011). Thus, malfunction of the ETC in mitochondria is usually accompanied with the loss of mitochondrial membrane potential (van der Stel et al., 2022). We performed TMRM staining to investigate if there is a link between the reduction OCR and a rupture of mitochondrial membrane potential caused by JNK inhibition. With TMRM staining, we could indeed observe a reduction in mitochondrial membrane potential upon JNK inhibition. Overall, there is multiple evidence that mitochondrial dysfunction is increased by JNK pathway inhibition in UdRPCs.

JNK signaling is involved in proliferation processes, consequently this pathway also regulates cellular respiration. Xie, Sun *et al*. demonstrated that inhibiting the JNK signaling pathway in hematopoietic stem cells results in a reduction in the expression of genes related oxidative phosphorylation (Xie et al., 2022). Our data indicates that inhibition of JNK signaling pathway leads to mitochondria impairment and ferroptosis. One of the hallmarks for iron-dependent cell death include morphological changes of the mitochondria such as a blistered cell membrane, reduction in size and loss of mitochondria cristae (Galluzzi et al., 2018; Li et al., 2020). Since mitochondrial complexes involved in oxidative phosphorylation are localized in the inner mitochondrial membrane, ferroptosis-related damage of mitochondria may be an explanation for the rupture of mitochondrial membrane and thus to an impaired oxidative phosphorylation (Nolfi-Donegan et al., 2020). Moreover, mitochondrial dysfunction via mitochondrial ROS production activates mitochondrial JNK signalling, which promotes Bax-dependent apoptosis (Chambers and LoGrasso, 2011; Niizuma et al., 2010). Based on this finding, JNK inhibition in our study could lead to an inhibition of Bax-dependent apoptosis, which simultaneously enhances ferroptosis.

## Conclusion

In this study, we demonstrated the importance of JNK signaling for the maintenance of self-renewal and the proliferation capacity SIX2-positive in urine-derived renal progenitor cells. Pathway inhibition led to the emergence of ferroptosis-induced cell death in UdRPCs and was accompanied by disrupted mitochondrial membrane potential and overall reduced oxidative phosphorylation. Therefore, we propose the use of JNK-inhibited UdRPCs as model for ferroptosis-induced kidney diseases such as acute kidney injury.

## Acknowledgements

J.A. acknowledges the medical faculty of Heinrich Heine University for financial support.

J.A., C.B., and A.S.R. acknowledge that this work is partly funded by the Deutsche Forschungsgemeinschaft (DFG, German Research Foundation) – 417677437/GRK2578.

## Author Contributions

L.N. designed and performed experiments, analysed the data, wrote and edited the manuscript. M.W. and L.T. assisted in experimental design, performed experiments, analysed data, wrote and edited the manuscript. W.W. prepared the formal analysis, data curation, helped with the figures and edited the manuscript. A.S.R. supervised oxidative phosphorylation-related experiments and edited the manuscript. C.B. supervised ferroptosis-related experiments and edited the manuscript. J.A. conceptualized the work, wrote and edited the manuscript, had the project administration, acquired funding and supervised the study. All authors have read and agreed to the published version of the manuscript.

## Declaration of interests

The authors declare no competing interests.

### LEAD CONTACT AND MATERIALS AVAILABILITY

Further information and requests for resources and reagents should be directed to and will be fulfilled by the Lead Contact, James Adjaye.

## Methods

### Cell Culture

For this study, three distinct UdRPC lines UM51, UM27 and UF21 were used. Their names describe the donor’s gender (UM-urine male; UF-urine female) and age. The cells were cultivated on 0.2 % gelatine-coated plates and were maintained in proliferation medium supplemented with 5 ng/ml bFGF (Peprotech) every second day (Rahman et al., 2020). The optimal concentration of the JNK inhibitor AEG3482 (TOCRIS) was determined by titration of different concentrations on UM51 cells. The concentrations 10 µM, 50 µM and 100 µM AEG3482 inhibitor were applied to UM51 cells for 48 h to 120 h with daily medium change. A concentration of 10 µM AEG3482 was adopted for subsequent experiments. At 80 % confluency, cells were trypsinized with TrypLE (Life Technologies) and were seeded on gelatine-coated plates. The treatment with 10 µM AEG3482 inhibitor was started at 60-70 % confluency and was maintained for 24 h, 72 h and 120 h. In parallel, untreated cells were kept as control for the same time points and were cultivated in proliferation medium with daily changes of medium. Like UM51 cells, the cell lines UM27 and UF21 were treated for 72 h with 10 µM JNK inhibitor AEG3482 with daily medium changes.

### Proliferation assay

After JNK inhibition, cells were fixed with 4 % paraformaldehyde (Polysciences) and stained with the antibody anti-mouse KI67, 1:200 (CST). Randomly chosen pictures were taken from each well and KI67-positive cells as well as total cell numbers were counted (N=10). The ratio of KI67-positive/total cell number was calculated and statistical analysis was processed in Graphpad Prism software (Dotmatics). *P*-values were calculated with an unpaired t-test (two-tailed). (* ρ-value < 0.05, ** ρ-value < 0.01, *** ρ-value < 0.001).

### Immunofluorescence

Cells were fixed with 4 % paraformaldehyde and were subsequently permeabilized with 0.5 % Triton X-100/PBS (Sigma-Aldrich) for 15 min. Prior to incubation with the primary antibodies, blocking with 3 % BSA (Sigma-Aldrich) for 1 h at room temperature was performed. Primary antibodies were diluted as following: anti-mouse KI67 (1:200), anti-rabbit cJUN (1:400) (CST), anti-rabbit phospho-cJUN (1:800) (CST). The plates were then incubated at 4°C overnight. Labelled cells were detected with the secondary antibodies-anti-rabbit Alexa Fluor™ 488 (1:500) (Thermofisher) and anti-mouse Alexa Fluor™ 555 (1:500) (Thermofisher). Nuclei were stained with Hoechst (1:5000) (Thermofisher). Pictures were taken under fluorescence microscope (LSM700; Zeiss) and processed with ZenBlue 2012 Software Version 1.1.2.0 (Zeiss) and Image J (NIH).

### Western Blotting

Total protein was extracted by lysing the cells with RIPA Buffer (Sigma-Aldrich) containing phosphatase and protease inhibitors (Sigma-Aldrich). Protein was quantified with the Pierce’s BCA assay kit from Thermo Scientific. Electrophoresis was run with a protein input of 20 µg. Proteins were bound with antibodies including anti-rabbit phospho-SAPK/JNK (1:1000) (CST), anti-rabbit SAPK/JNK (1:1000) (CST), anti-rabbit cJUN (1:1000) (CST), anti-rabbit phospho-cJUN (1:1000) (CST), anti-rabbit SMAD2/3 (1:1000) (CST), anti-rabbit phospho-SMAD2 (CST) (1:1000), anti-rabbit SMAD1 (1:1000) (CST), anti-rabbit phospho-SMAD1/5 (1:1000) (CST), anti-rabbit GPX4 (1:1000) (CST) and anti-mouse β-actin (1:5000) (CST). Enhanced chemiluminescent (ECL) horseradish-peroxidase (HRP) detection technique was used to detect the specific proteins (Life Technologies).

### Flow cytometry-based measurement of lipid peroxidation

Untreated and JNK-inhibited cells were cultivated for 72 h with daily medium change. For inhibition of ferroptosis, cells were additionally treated with 100 nM Liproxstatin-1 (Lip-1) (Sigma-Aldrich) 1h before JNK inhibition via AEG3482. Cells were washed twice with PBS and stained with 1 µM BODIPY™ 581/591 C11 (Invitrogen) for 15 min at 37 °C and 5 % CO2. Subsequently, cells were washed twice with PBS and harvested with trypsin. The samples were centrifuged for 10 minutes at 700 g and resuspended in 300 µl MACS buffer (0.5 % BSA, 2 mM EDTA, PBS). Lipid peroxidation was detected by measuring the fluorescence intensity of BODIPY 581/591 C11 using BD FACS Canto 2 (BD Biosciences). In each case, 3×10^4^ events were measured, and the data evaluated using FlowJo software (BD Biosciences). Statistical significance was calculated according to unpaired t-test (two-tailed) using GraphPad Prism software (n=5; * ρ-value < 0.05, ** ρ-value < 0.01, *** ρ-value < 0.001).

### Seahorse XF Cell Mito Stress Test

Optimal seeding density and FCCP concentration without toxicity response were tested before starting the Seahorse XF Cell Mito Stress Test assay (Agilent). A cell density of 4×10^3^ cells per well and a FCCP concentration of 2 µM was determined and used for the following assays. Cells of the three UdRPC lines UM51, UM27 and UF21 were seeded on gelatine-coated Seahorse XF Cell Culture Microplates. For mitochondrial oxygen consumption rate (OCR) measurements of JNK pathway inhibition, cells were treated with or without JNK inhibitor AEG3482 for 72 h. Following the manufacturer’s protocol, XF sensor cartridges were hydrated with Seahorse XF Calibrant at 37°C in a hypoxic incubator overnight. The medium was changed to phenol-free Seahorse XF DMEM (10 mM glucose, 1mM pyruvate and 2 mM L-glutamine (all from Sigma Aldrich). The OCR was measured in a Seahorse XFe96 Flux Analyser with Seahorse Wave 2.4 software (Agilent). In 3 cycles of 3 min mixing and 3 min recording, 1 µM Oligomycin, 2 µM FCCP and 0.5 µM Antimycin/Rotenone AA was injected and OCR was measured. The OCR of UF21, UM51 and UM27 was measured with one biological replicate (UM51 and UM27: N=23; UF21: N=46). The data was normalized to the cell number and statistical significance was calculated with an unpaired t-test (* ρ-value < 0.05, ** ρ-value < 0.01, *** ρ-value < 0.001).

### TMRM Staining

Cells were seeded on MatTek glass bottom dishes for microscopy followed by a cultivation time of 72 h with or without AEG3482 and daily media change. Cells were then stained with Tetramethylrhodamin (TMRM) and MitoTracker™Green FM (Invitrogen) for 30 min. Briefly, cells were washed thrice with 1 x PBS and were covered in Opti-MEM™(Gibco) without phenol red containing 10 µM HEPES buffer for buffering oxidation of media. Fluorescence imaging of UdRPCs was acquired at Nikon Ti2 inverted confocal microscope, coupled with UltraVIEW®VoX spinning disc laser system (PerkinElmer) equipped with a 63-x oil objective (N.A. 1.2). Imaging was performed in a chamber at 37 °C. For analysis of the fluorescence images, background correction was performed by manually defining a region of interest (ROI) in the background of the image and subtraction of the mean fluorescence signal in all images. For each individual cell, a ROI was manually defined for measurement of TMRM fluorescence intensity. Background correction and TMRM intensity measurement of 50 cells per sample was performed with Volocity® Software for spinning disk microscopy. High variations in fluorescence intensities were cleaned by an outlier test ROUT (Q = 1%). Cleaned data was used for statistical and graphical analysis, performed in GraphPad Prism software. Statistical significance was calculated with an unpaired t-test (n=2; control: N= 1; JNK inhibition: N= 2; * ρ-value < 0.05, ** ρ-value < 0.01, *** ρ-value < 0.001).

### Quantitative RT-PCR

RNA was isolated with the DIrectzol RNA MiniPrep kit (Biozol) according to the manufacturer’s protocol. Subsequently, 500 ng RNA was transcribed to cDNA using MultiScribe Reverse Transcriptase (Life Technologies). Quantitative PCRs were performed in technical triplicates using POWER SYBR Green Master Mix (Life Technologies). Primer sequences are listed in *Table S1* (purchased from MWG). Mean values were normalized to the housekeeping gene *RPL37A*, compared to an untreated control for the specific time point and calculated by the 2 −ΔΔCt method.

### Gene expression analysis

Total RNA of UdRPCs treated with the JNK inhibitor AEG3482 and untreated cells was hybridized onto microarrays of type Affymetrix Human Clariom S assay at the BMFZ (Biomedizinisches Forschungszentrum) core facility of the Heinrich-Heine University, Düsseldorf. The R/Bioconductor environment (Gentleman et al., 2004) was employed to process the Affymetrix microarray data. The data was background-corrected via the Bioconductor package oligo (Carvalho and Irizarry, 2010) and normalized applying the Robust Multi-array Average (RMA) method. The packages VennDiagram (Chen and Boutros, 2011) and gplots (Warnes et al., 2005) were used to generate Venn diagrams of the numbers of genes expressed in the control or JNK-inhibition conditions. A gene was considered expressed when the detection p-value -determined as described in Nguyen *et al*. (2022) was below the threshold of 0.05 (Nguyen et al., 2022). Hierarchical clustering was analyzed via (i) the R function *hclust* parametrized with Pearson correlation as similarity measure and complete linkage as agglomeration method in dendrograms, (ii) the R function *heatmap*.*2* from the gplots package (Warnes et al., 2005) and also parametrized with Pearson correlation as similarity measure and additionally with colour-scaling per row/gene in heatmaps.

### Gene ontology (GO) and pathway analysis

Upregulated genes were calculated by the criteria: ratio between the JNK-inhibited state and control greater than 1.5 and detection-p-value in the JNK-inhibited state below the threshold of 0.05, down-regulated genes analogously by the criteria: ratio between the JNK-inhibited state and control less than 0.67 and detection-p-value in the control state below the threshold of 0.05. From the resulting gene sets over-represented GOs were determined via the Bioconductor package GOstats (Falcon and Gentleman, 2007). KEGG (Kyoto Encyclopedia of Genes and Genomes) (Kanehisa et al., 2017) pathways were analyzed for over-representation based on associations between pathways and genes downloaded from the KEGG database in July 2020. For each of the KEGG pathways the hypergeometric test was applied to the sets of up- and downregulated genes, which could be annotated to that pathway. Furthermore, genes from KEGG pathways as well as gene sets found by single cell sequencing analysis of fetal kidney development by Lindström *et al*. (2018) were employed to generate heatmaps for cluster analysis as described above (Lindström et al., 2018).

### Metascape Analysis

Gene enrichment analyses of differential GO/KEGG terms and biological processes between JNK-inhibited UdRPCs and untreated controls were performed using metascape (http://metascape.org; (Zhou et al., 2019)). Exclusive gene sets of JNK inhibition and control of each time point (24 h, 72 h, 120 h) were used as data source. The metascape software applied hierarchical clustering to display calculated significant GO terms into a tree, which was spread into term clusters with a 0.3 kappa score as a threshold. The top enrichment clusters were represented as heatmaps with a color scale ranging from gray to dark orange. Statistical significance was hereby displayed in dark orange and lack of enrichment in gray color.

### Data Availability

Microarray raw data have been deposited at NCBI GEO and are publicly available as of the date of publication. Accession number is listed in the key resources table. All data reported in this paper will be shared by the lead contact upon request.

Any additional information required to reanalyse the data reported in this paper is available from the lead contact upon request.

## Supplemental Information

**Figure S1. Inhibition of JNK and FGF induces stress in UdRPCs (Related to Figure 1 and 4)**. A) Morphological analysis of cellular stress and cell death in UM51 treated with 50 µM and 100 µM AEG3482 inhibitor. Scale bars represent 100 µm. B) Morphological analysis of cellular stress and cell death in UF21 treated with 50 µM and 100 µM AEG3482 inhibitor. Scale bars represent 100 µm. C) Representative histograms of measured fluorescence intensities after BODIPY staining and the respective bar plots of mean fluorescence intensity of control, JNK inhibition and JNK inhibition+Lip-1 (n=5; * ρ-value < 0.05, ** ρ-value < 0.01, *** ρ-value < 0.001). Error bars indicate SEM. D) Representative histograms of measured fluorescence intensities after BODIPY staining and the respective bar plots of mean fluorescence intensity of control or FGF-inhibited UdRPCs (n=5; * ρ-value < 0.05, ** ρ-value < 0.01, *** ρ-value < 0.001). Error bars indicate SEM.

**Table S1: List of qRT-PCR primers (Related to Figure 1 and 4)**.

**Table S2: Lists of exclusive and common genes of the Venn analysis for 24h (Related to Figure 2)**.

**Table S3: Lists of exclusive and common genes of the Venn analysis for 72h (Related to Figure 2)**.

**Table S4: Lists of exclusive and common genes of the Venn analysis for 120h (Related to Figure 2)**.

**Table S5: GO terms of down- and up-regulated genes from KEGG analysis for the time points 24 h, 72 h and 120 h after JNK inhibition (Related to Figure 2)**.

**Table S6: Downregulated cell cycle-related gene lists of Metascape analysis for the time points 24 h, 72 h and 120 h after JNK inhibition (Related to Figure 2)**.

