## Supplementary figures and images for "JNK signalling regulates self-renewal of proliferative urine-derived renal progenitor cells via inhibition of ferroptosis"

### Supplemental Figure 1

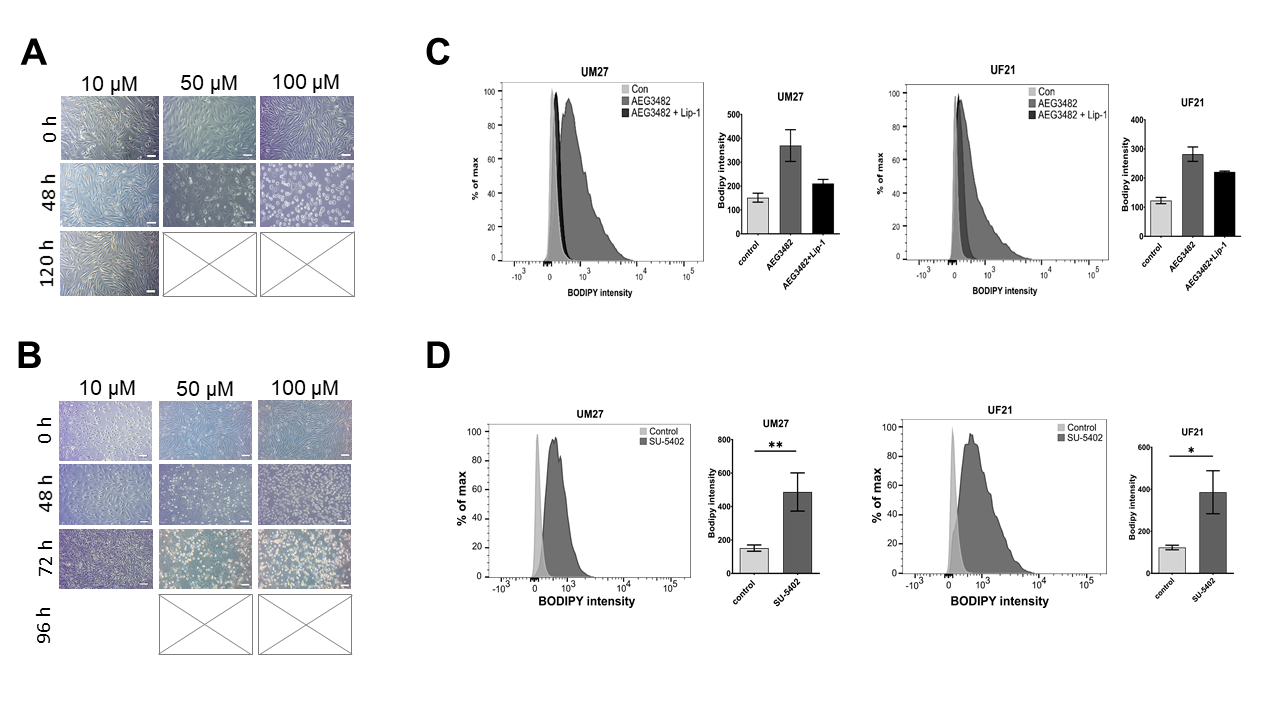
